# Multi-scale structural similarity embedding search across entire proteomes

**DOI:** 10.1101/2025.02.28.640875

**Authors:** Joan Segura, Ruben Sanchez-Garcia, Sebastian Bittrich, Yana Rose, Stephen K. Burley, Jose M. Duarte

## Abstract

The rapid expansion of three-dimensional (3D) biomolecular structure information, driven by breakthroughs in artificial intelligence/deep learning (AI/DL)-based structure predictions, has created an urgent need for scalable and efficient structure similarity search methods. Traditional alignment-based approaches, such as structural superposition tools, are computationally expensive and challenging to scale with the vast number of available macromolecular structures. Herein, we present a scalable structure similarity search strategy designed to navigate extensive repositories of experimentally determined structures and computed structure models predicted using AI/DL methods. Our approach leverages protein language models and a deep neural network architecture to transform 3D structures into fixed-length vectors, enabling efficient large-scale comparisons. Although trained to predict TM-scores between single-domain structures, our model generalizes beyond the domain level, accurately identifying 3D similarity for full-length polypeptide chains and multimeric assemblies. By integrating vector databases, our method facilitates efficient large-scale structure retrieval, addressing the growing challenges posed by the expanding volume of 3D biostructure information.

## Introduction

Recent advances in deep neural architectures have revolutionized large-scale prediction of 3D structures of biological macromolecules. Deep learning models (Baek *et al*., 2021; Du *et al*., 2021), most notably AlphaFold2 (Jumper *et al*., 2021; Abramson *et al*., 2024), have demonstrated remarkable success in predicting protein structures with accuracies comparable to that of lower-resolution experimentally determined structures (Shao *et al*., 2022), significantly accelerating structural biology research (Yang *et al*., 2023; Terwilliger *et al*., 2024). More recently, models like the Evolutionary Scale Modeling 3 (ESM3) (Hayes *et al*., 2025) have further expanded the horizon of applications by leveraging large-scale protein language models (PLM) to predict both structure and function. These manifold breakthroughs have led to the creation of extensive structural databases, such as AlphaFold DB (Varadi *et al*., 2024), which now hosts more than 214 million structures predicted by Google Deep Mind, ESMAtlas (Lin *et* al., 2023), a database of over 600 million protein structures derived from metagenomic sequences, and the ModelArchive (Tauriello *et al*., 2025), a repository for computed structure models open to all contributors. By providing high-quality structural data at an unprecedented scale, these advances are transforming protein structure analysis and reshaping computational approaches to process structural information efficiently.

The Protein Data Bank (PDB) (Berman *et al*., 2003) serves as the global repository for experimentally determined macromolecular structures, providing an essential resource for structural biology, structure-guided drug discovery, and structural bioinformatics. Jointly managed by the Research Collaboratory for Structural Bioinformatics Protein Data Bank (RCSB PDB, RCSB.org) in the United States, together with its Worldwide Protein Data Bank (wwPDB, wwpdb.org) partners, the PDB has long been the gold standard for high-resolution 3D biostructure data generated by macromolecular crystallography, 3D electron microscopy, and nuclear magnetic resonance spectroscopy. Importantly, the success of deep learning-based structure prediction methods—such as AlphaFold2, RoseTTAFold, and ESM3—would not have been possible without the vast repository of rigorously validated, expertly biocurated experimental structures stored in the PDB. PDB data were essential for training these and other AI/DL models. Recognizing the growing impact of deep learning prediction methods, RCSB.org has expanded its research-focused web portal to deliver computed structure models (CSMs) (Burley *et al*., 2023), including those generated by AlphaFold2 and those deposited into the ModelArchive. Currently, RCSB.org hosts >1 million CSMs, providing open access to the entire proteomes of human, model organisms, agriculturally important plants, select bacterial pathogens, and species relevant to understanding and mitigating climate change (Burley *et al*., 2025). By incorporating these predicted structures alongside experimentally determined structures, RCSB.org provides many millions of researchers, educators, and students with a comprehensive one-stop shop for 3D biostructural analysis, functional annotation, and drug discovery. RCSB.org bridges the gap between experimentally validated data and large-scale predictive modeling, facilitating new avenues for understanding protein function, evolution, and biologically important molecular interactions at unprecedented scale.

As the volume of available 3D biostructure data continues to grow, efficient searching and comparisons among structures at scale have become critical challenges for computational structural biologists. With resources such as AlphaFold DB and the ModelArchive hosting hundreds of millions of predicted protein structures, and RCSB.org expanding its catalog of integrated CSMs, structural similarity search methods must support real-time, accurate queries across a vast and continuously growing 3D biostructure space. Moreover, structure comparisons are inherently complex due to the multi-scale nature of macromolecular architecture (spanning primary, secondary, tertiary, and quaternary levels). Macromolecular structures can be analyzed at different hierarchical levels of granularity, including individual domains, full-length protein chains, and larger multimeric assemblies, each of which may exhibit unique structural and functional properties. Effective search algorithms must account for these varying levels of granularity while maintaining computational efficiency, ensuring that relevant structural relationships can be identified across diverse datasets. Developing scalable and precise structural similarity search methods is, therefore, essential for unlocking new biological insights and enabling the functional annotation of uncharacterized proteins and assemblies within the rapidly expanding corpus of structural information.

Traditional structure superposition tools (Zhang *et al*., 2022; Holm, 2019) are computationally complex, rendering them unsuitable for large-scale, real-time similarity searches. To improve search performance, bioinformaticians have relied on large-scale computational resources to handle computationally intensive steps (Krissinel and Henrick, 2004) or previous clustering based on structure to reduce the search space (Prlic *et al*., 2010). Both such approaches have limited scalability.

Recently developed methods offer various solutions for searching large volumes of structural data while aiming to retain high levels of sensitivity. One such approach, BioZernike descriptors (Guzenko *et al*., 2020), encodes macromolecular structures as vectors of features using Zernike polynomials and global geometric measures. A logistic regression model, trained using the CATH classification as ground truth, is then applied to compare pairs of descriptors. Although this method efficiently handles the current volume of experimental structural data, its sensitivity is lower compared to other approaches.

A complementary approach, Foldseek (van Kempen *et al*., 2024), transforms 3D protein structures into sequences using a structural alphabet, effectively reducing the structural search problem to a sequence similarity search task. When integrated with the sequence search software MMseqs (Steinegger and Söding, 2017), Foldseek enables real-time searches across large datasets, provided that the data is clustered and sufficient computational resources are available. Currently, Foldseek is employed as the structural search method in the AlphaFold DB server, utilizing a pre-clustered subset of the 214 million structures at 50% sequence identity to improve performance.

Embedding-based approaches transform complex data such as text, audio, and images into high-dimensional vector representations that capture meaningful relationships in a continuous space. Such embeddings enable efficient similarity searches by positioning similar entities closer together in the vector space, allowing for fast and scalable comparisons. To manage the growing volume of high-dimensional data, vector databases store and index these embeddings, enabling large-scale retrieval across millions of entries by employing Approximate Nearest Neighbors (ANN) algorithms. This approach has revolutionized multiple fields, including natural language processing (Xu *et al*., 2023) and computer vision (Covington *et al*., 2016). In computational biology, the TM-Vec approach applies this concept to protein sequences, encoding them as vectors to facilitate remote homology detection (Hamamsy *et al*., 2024). Herein, we present a scalable structural similarity search strategy designed to traverse enormous volumes of 3D biostructure data, including both experimentally determined atomic coordinates and CSMs. Our approach follows an embedding-based methodology, transforming macromolecular structures into fixed-length vectors and leveraging vector databases for optimal search performance. The method integrates the ESM3 generative model to convert 3D structures into sequences of residue embeddings, which are then processed by a transformer-based network that aggregates the information into a fixed-length vector. The embedding model was trained to predict TM-scores between pairs of 3D structures. Despite being trained exclusively on single-domain structures, our results demonstrate that the model generalizes beyond the domain level to full-length polypeptide chains (frequently encompassing multiple structurally dissimilar domains). Moreover, we extended the embedding representation to macromolecular assemblies, achieving competitive performance in structure similarity searches for multi-protein complexes. Finally, we stood up a proof-of-concept (PoC) system that stores embeddings for 214 million CSMs from AlphaFold DB in a vector database, showing the immediate feasibility of handling the current structural knowledge with modest computational resources. The PoC system is publicly available at http://embedding-search.rcsb.org, allowing queries for PDB and AlphaFold DB entries or custom user-uploaded files.

## 2 Materials and Methods

### 2.1 Datasets

This section introduces the various datasets used for training and testing our embedding model.

Domain pairs training set: The training set consists of 15,176 SCOPe40 (v2.08)(Chandonia *et* al., 2022) domains from the ASTRAL subset, a non-redundant collection with <40% sequence identity, corresponding to protein domains identified in PDB structures. These domains represent 4,703 families and have an average length of 182 amino acids. The dataset was compiled by Chengxin Zhang, and all associated information, including domain coordinate files, is publicly available in the Zenodo repository for research purposes (Zhang, 2022). Both TM-scores, query, and target, are available for all domain pairs, resulting in over 115 million comparisons. For training, we used the highest TM-score in each pair. This dataset of domain pairs and TM-scores will be referred to as DT115M hereafter.

Domain pairs benchmarking: To evaluate the performance of the embedding model for structural similarity search and compare it to results obtained with Foldseek and other structural search methods, we used the benchmark described in reference (van Kempen *et al*., 2024). This dataset consists of 11,211 single domains from SCOPe40 (v2.01), with an average length of 174 residues, representing 4,161 distinct families (hereafter DS62M). All domain pairs (>62 million) in the dataset were used to assess the capability of each method to identify matches at the family, superfamily, and fold levels.

Full-length chain benchmarking: A non-redundant set of full-polypeptide chain protein structures was harvested from the PDB. We used RCSB.org sequence clusters to guarantee an identity level of less than 30% between proteins. Subsequently, only proteins with sequence lengths longer than 200 amino acids and fewer than 20 structurally unmodelled residues were considered for the final collection resulting in 7,899 protein chains. Finally, we computed the TM-score value for all possible pairs using the US-align software enabling the ‘-fast’ configuration option. It should be noted that TM-score may underrepresent similarity in multi-domain proteins where domain movements occur, although such cases are relatively uncommon among PDB entries (Burra *et al*., 2009). The resulting dataset (hereafter PS31M) contained >31 million protein pair comparisons. This dataset was used to evaluate the performance of our approach for structural similarity searches in full-polypeptide chain protein cases longer than the average single domain lengths used in the other benchmarks.

Predicted structures dataset: We compiled a dataset of CSMs derived from the non-redundant AlphaFold DB set used in previous Foldseek benchmarks (van Kempen *et al*., 2024). From the initial collection of >34,000 proteins, we applied additional filtering based on domain content using The Encyclopedia of Domains (TED) (Lau *et al*., 2024) and the CATH classification (Waman *et al*., 2024). Only proteins containing two or more high-confidence domains assigned in CATH were retained, yielding a final dataset of 2,155 multidomain proteins spanning 1,038 distinct domain families. All possible domain pair combinations from this dataset (hereafter AF23M) were used to evaluate the ability of different methods to identify proteins sharing domains within the CATH classification.

Assembly Dataset: To evaluate the performance of our embedding model for structural similarity searches in multimeric protein assemblies, we selected, at random, 10,000 homomeric assemblies from the 238,965 structures in the 3D Complex database V7.0 (Levy *et al*., 2006). For each possible pair, we computed TM-scores using the US-align tool with the ‘-fast’ configuration. The resulting dataset (hereafter AS50M) contained over 50 million assembly pairs with annotated TM-scores.

### 2.2 Embedding Model

The embedding model transforms protein structures into numerical vectors of fixed length. The model consists of two main components: (1) a PLM that computes an embedding for each residue in a given 3D structure, and (2) a transformer-based neural network that aggregates these residue-level embeddings into a single vector, encoding information about the entire 3D structure. Figure 1 displays a visual representation of the embedding model and its different components.

**Figure 1.**
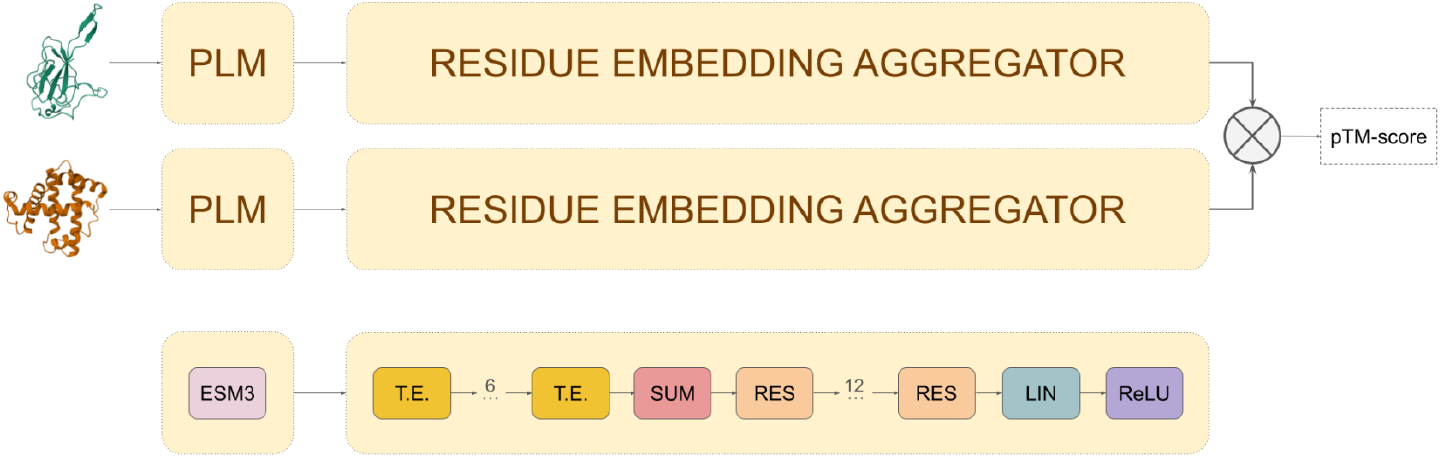
Embedding model network architecture. The model operates as a twin neural network, utilizing shared weights to calculate embeddings for pairs of 3D structures and predict TM-scores from their cosine distance. The model consists of two main components: (1) a protein language model (PLM), ESM3, which encodes protein 3D structures as residue-level embeddings, and (2) an aggregator module that processes these embeddings into a fixed-length vector. The aggregator comprises six transformer encoder layers (T.E.), a summation pooling operation (SUM), and twelve fully connected residual blocks (RES).

Residue-level embeddings are computed using ESM3, a generative PLM (Hayes *et al*., 2025). Given the 3D structure of a single protein chain, this model outputs a 1,536-dimensional vector for each residue, capturing both sequence and structural information.

The aggregation network processes the residue-level embeddings from ESM3 and outputs a fixed-length vector. This component consists of six stacked transformer encoders (Vaswani *et* al., 2017), each with feedforward layers containing 3,072 neurons and ReLU activations. Following the transformer encoders, a summation pooling operation and 12 fully connected residual layers (He *et al*., 2016) of 1,536 neurons aggregate the resulting embeddings into a single 1,536-dimensional vector, encoding information for the entire 3D structure. The aggregator module contains a total of ∼170 million trainable parameters.

### 2.3 Model Training

Our embedding model was trained to predict the maximum TM-score between pairs of 3D structures. TM-score values were rounded to the first decimal place allowing the network to approximate 11 possible TM-score values:

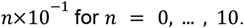

During training, the model operated as a twin neural network, utilizing shared weights to produce embeddings for pairs of 3D structures (see Figure 1). The ESM3 model weights are frozen and the aggregator network parameters were optimized to minimize the mean square error between the cosine similarity of the computed embeddings and the TM-score of their superposed 3D structures.

The final model was trained using domain pair batches from the DT115M dataset. Batches were randomly generated ensuring that the TM-score values of the domain pairs were uniformly distributed among the 11 possible values. This strategy addresses the unbalanced nature of TM-score values within the DT115M dataset, where only 0.05% of TM-score values exceed 0.8 and more than 95% fall under 0.4 (see Supplementary Material Figure S1).

To evaluate model performance and store checkpoints during the training process, a validation dataset was created by removing 2% of the DT115M training dataset. After each epoch, the area under the precision-recall curve (AUPRC) was calculated on the validation set as a measure of performance. The model was trained for 100 epochs, each consisting of 320,000 randomly selected domain pairs. The best performance was achieved at epoch 31, with no significant improvement thereafter. We used the Adam optimizer (Kingma and Ba, 2015) with a learning rate schedule incorporating cosine decay and a two-epoch warmup. Training was conducted using 64 A100 GPUs.

### 2.4 Assembly embeddings

We extended our model to predict embeddings for multimeric assemblies using the following strategy. For a given assembly, the generative model ESM3 was used to compute residue-level embeddings for each chain individually. Since ESM3 operates at the chain level, the resulting per-residue embeddings are independent of the chain order in the assembly. Then, chain residue-level embeddings were concatenated to form a single sequence of embeddings. Subsequently, our residue-level aggregator model was applied to compute a fixed-length vector representation for the whole 3D structure of the assembly. Since the aggregator module does not include any positional encoding, it remains equivariant to chain permutations, and when combined with the summation pooling operation, this ensures that the final embedding representation is invariant to the selected chain order in the assembly.

Although our model was trained exclusively on single domains, Section 3.4 demonstrates how the latent features learned by our embedding model generalize beyond single-domain structures, enabling it to perform structural similarity searches at the assembly scale.

### 2.5 Structural Embedding Database

A dedicated vector database (Milvus) was deployed to store embeddings for all structures integrated into RCSB.org, including both experimental entries and CSMs. More than 2 million embeddings were computed, covering all individual chains and biological assemblies, and subsequently indexed using the Hierarchical Navigable Small World (HNSW) algorithm for ANN searches(Malkov and Yashunin, 2020). The HNSW algorithm constructs multi-layered graphs that efficiently balance speed and accuracy, significantly reducing search complexity for large-scale vector datasets.

To demonstrate the scalability of the embedding approach, we generated embeddings for all predicted structures (>214 million) available in AlphaFold DB (Varadi *et al*., 2024) and stored them in a separate vector database. Due to the large data volume, the DiskANN indexing approach(Jayaram Subramanya *et al*., 2019) was used to create a disk-based index while maintaining efficient memory use and search speed.

## 3. Results

### 3.1 Protein domain benchmark

Our primary objective in developing this embedding model is to identify 3D structures similar to a user-provided query. To evaluate its performance and to compare it with Foldseek and other state-of-the-art methods^20^, we employed the DS62M benchmark dataset. Two complementary evaluation strategies were applied: (i) an all-versus-all domain comparison, where sensitivity was measured as the fraction of true positives (TPs) retrieved before the first false positive (FP), and (ii) precision–recall analysis, with the area under the curve (AUC) providing a global performance metric.

Performance was assessed at three hierarchical levels of structural classification: domain family, superfamily, and fold. TPs were defined as pairs belonging to the same family, pairs belonging to the same superfamily but different families, and pairs sharing the same fold but different superfamilies, respectively. Domains from different folds were classified as FPs. Results at the domain family level are presented in Figure 2, while superfamily- and fold-level results are provided in the Supplementary Material (Figure S2).

**Figure 2.**
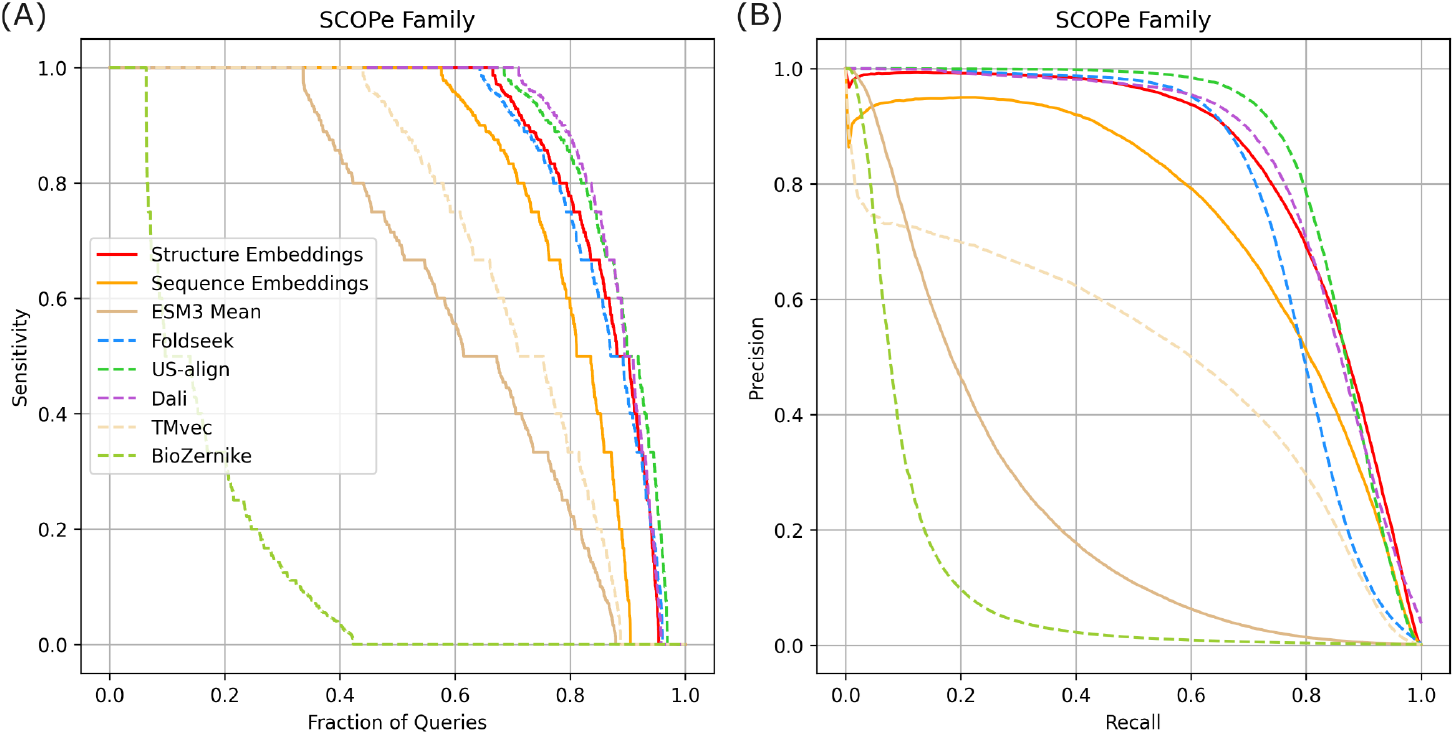
Sensitivity and precision-recall performance on SCOPe40 v2.01 domain structures. Each domain was compared against all others in the dataset, and results were ranked in descending order of predicted similarity score. True positives (TPs) were defined as matches within the same family, and false positives (FPs) as matches between different folds. (A) Sensitivity curves across methods. Sensitivity for each domain was calculated as the number of TPs retrieved before the first FP; values are shown sorted from best to worst as a function of the fraction of queries. (B) Precision–recall curves across methods.

Figure 2A shows the distribution of sensitivities across queries, sorted from best to worst. Although the embedding model does not reach the sensitivity of computationally intensive alignment-based methods such as DALI and US-align, its performance is comparable to Foldseek and markedly superior to our earlier BioZernike descriptor approach. DALI and US-align achieved perfect sensitivity for 71% and 69% of domains, respectively, whereas our embedding model and Foldseek reached 66% and 64%. Precision–recall analysis (Figure 2B) further supports this trend: US-align and DALI obtained the highest AUC values (0.86 and 0.84, respectively), followed by our embedding model (0.83) and Foldseek (0.78). Complete AUC values for all methods are provided in Supplementary Material Table S1.

The benchmarks also included two sequence-based approaches: TM-vec (Hamamsy *et al*., 2024), a deep learning method for remote homology detection, and an alternative version of our model trained exclusively on sequence embeddings. Both approaches underperformed relative to our structurally trained embedding model (precision–recall AUC of 0.5 and 0.73, respectively). These results confirm that incorporating 3D structural information into residue-level embeddings yields substantially higher sensitivity and recall than sequence-only methods.

We additionally tested whether directly averaging residue embeddings from ESM3 could provide an effective representation of 3D structures. This naïve strategy resulted in markedly reduced performance (precision–recall AUC = 0.24), underscoring the necessity of the aggregator module for learning informative global embeddings.

Because SCOPe domain families in DS62M overlap with those in the DT115M training set, the benchmark described above employed a 10-fold cross-validation scheme to avoid redundancy between training and testing. The dataset was partitioned into 10 subsets, each excluding a unique collection of SCOPe families from training. Importantly, the training procedure was based solely on TM-score prediction between domain pairs, with no explicit use of SCOPe classification. This ensured that domains in the testing fold were unseen during training. However, as the benchmark involves all-versus-all comparisons, left-out queries were still compared against domains included in the training set. Supplementary Material Figure S3 illustrates this evaluation setup.

To ensure complete independence between training and testing, we conducted an additional control benchmark. Five percent of the unique domain superfamilies from DT115M were randomly withheld, and all domains belonging to these superfamilies were excluded from training. Sensitivity and precision–recall curves were then computed on the held-out domains. Supplementary Material Figure S4 and Table S2 present results from five such randomly sampled testing sets, showing performance consistent with the previous benchmark. Our embedding model achieved results comparable to US-align and Foldseek. These findings confirm that the model generalizes effectively to previously unseen domain families, successfully identifying structurally related pairs absent from the training data.

### 3.2 Full-length chain benchmark

Our previous benchmarks relied on domain-level datasets to assess the performance of the embedding model in identifying structural similarity. However, a more common use case involves retrieving structures similar to an entire protein chain. To evaluate whether our approach generalizes beyond the domain-level granularity used during training, we tested performance on the PS31M dataset, a non-redundant collection of full-length protein chains extracted from the PDB. For comparison, we also included other methods evaluated in Section 3.1.

Sensitivity and precision–recall analyses were conducted following a similar methodology as in Section 3.1. In this benchmark, TPs were defined using three different TM-score thresholds (0.6, 0.7, and 0.8), while FPs were protein pairs with TM-scores below 0.5 (proteins of distinct folds (Zhang and Skolnick, 2005)). Employing TM-scores as a baseline establishes a consistent reference for quantifying the divergence between the evaluated methods and US-align. Results for the strictest threshold (TM-score > 0.8) are shown in Figure 3, with analyses for the other thresholds available in Supplementary Material Figure S5 and Table S3.

**Figure 3.**
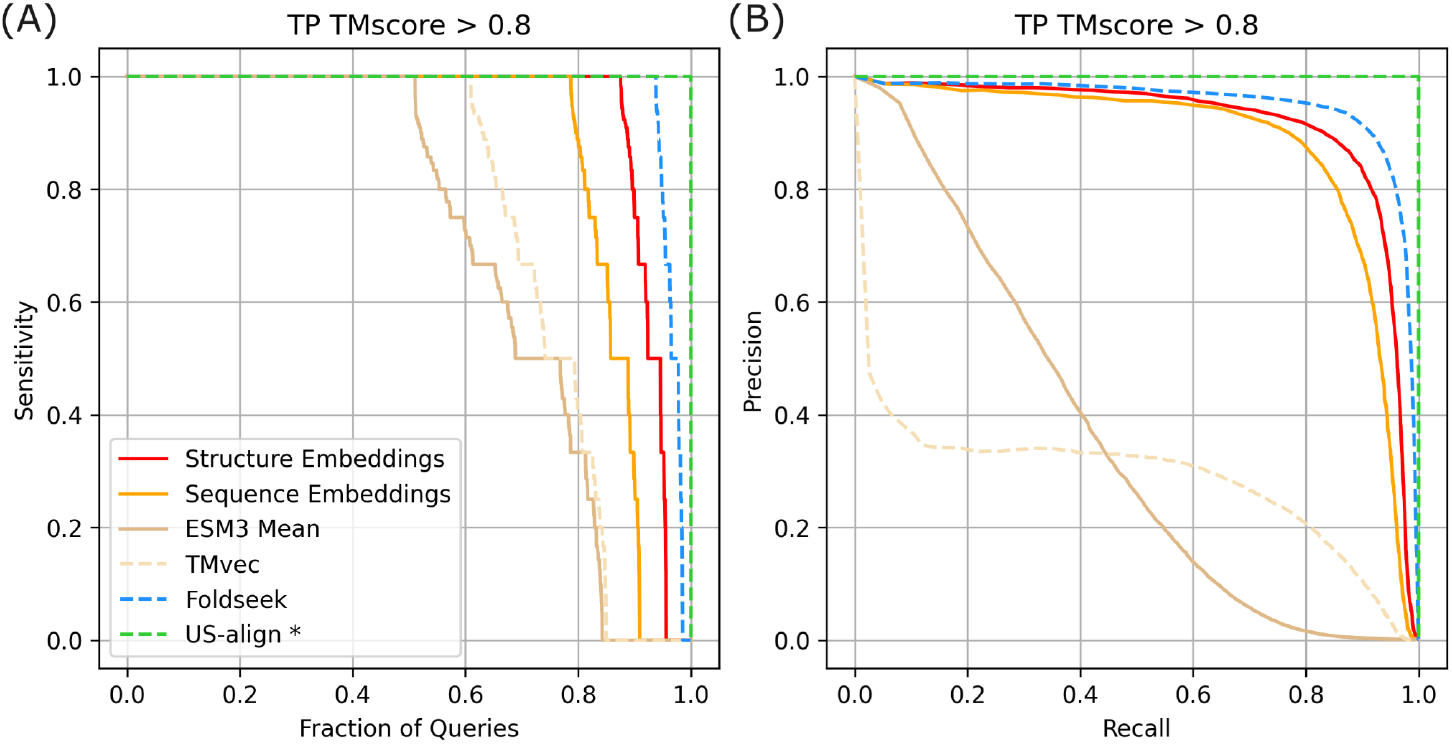
Sensitivity and precision-recall performance for a non-redundant set of PDB protein chain structures. Each protein chain was compared against all others in the dataset, and results were ranked in descending order of predicted similarity score. True positives (TPs) were defined as matches with TM-score > 0.8, and false positives (FPs) as matches with TM-score < 0.5. (A) Sensitivity curves across methods. Sensitivity for each chain was calculated as the number of TPs retrieved before the first FP; values are shown sorted from best to worst as a function of the fraction of queries. (B) Precision–recall curves across methods. US-align* shows perfect performance because it was used to compute the TM-score ground truth.

Figure 3A presents sensitivity to the first FP across methods. As the benchmark ground truth was defined by US-align, this method achieved a perfect classification. Foldseek achieved the highest performance, with 94% of queries reaching perfect sensitivity. Our embedding model closely followed, attaining 100% sensitivity for 89% of the queries. Precision–recall analysis (Figure 3B) revealed a similar trend: Foldseek reached an AUC of 0.95, while our embedding model achieved 0.91.

Consistent with the domain-level benchmark, sequence-based embedding approaches (TMvec and sequence-only embeddings) underperformed relative to embeddings that incorporate 3D structural information. Both approaches showed reduced sensitivity and recall compared with the structurally trained model. Our sequence-only embedding achieved higher performance than TMvec, reaching 100% sensitivity for 79% of queries and a precision–recall AUC of 0.88, compared with 61% sensitivity and an AUC of 0.29 for TMvec. Incorporating structural coordinates into residue embeddings substantially improved both sensitivity and recall in protein similarity searches.

Finally, direct averaging of ESM3 residue embeddings to encode full protein chains resulted in markedly reduced performance (precision–recall AUC = 0.35), highlighting the importance of the aggregator module in generating informative global embeddings.

Overall, this benchmark demonstrates that our embedding model extends beyond the original domain-based training set and generalizes effectively to full-length protein chains, enabling accurate structural similarity searches at the chain level.

### 3.3 Predicted structure models benchmark

A central motivation for our embedding approach is to provide a structural similarity search strategy that scales to the rapidly growing corpus of CSMs. To evaluate performance in this context, we used the AF23M dataset, which comprises multidomain proteins from AlphaFold DB. Although individual domain structures in CSMs are typically accurate, the relative positioning and orientation of domains are less reliable (Xia *et al*., 2023). As a result, approaches that rely solely on global structural similarity may fail to detect local domain-level similarities, making structural searches in these cases particularly challenging.

We assessed performance based on each method’s ability to identify protein pairs sharing the same domain composition at the CATH topology level. Following the strategy used in the protein domain benchmark (Section 3.1), TPs were defined as protein pairs with identical combinations of CATH topologies, whereas FPs were pairs in which all domains belonged to different CATH architectures. As in Section 3.1, sensitivity to the first FP and precision–recall curves were calculated after an all-versus-all comparison (Figure 4). Results at the CATH architecture level are provided in Supplementary Material Figure S6 and Table S4. Analysis at the homology level was not included, as many domains in the TED resource are classified only up to the topology level.

**Figure 4.**
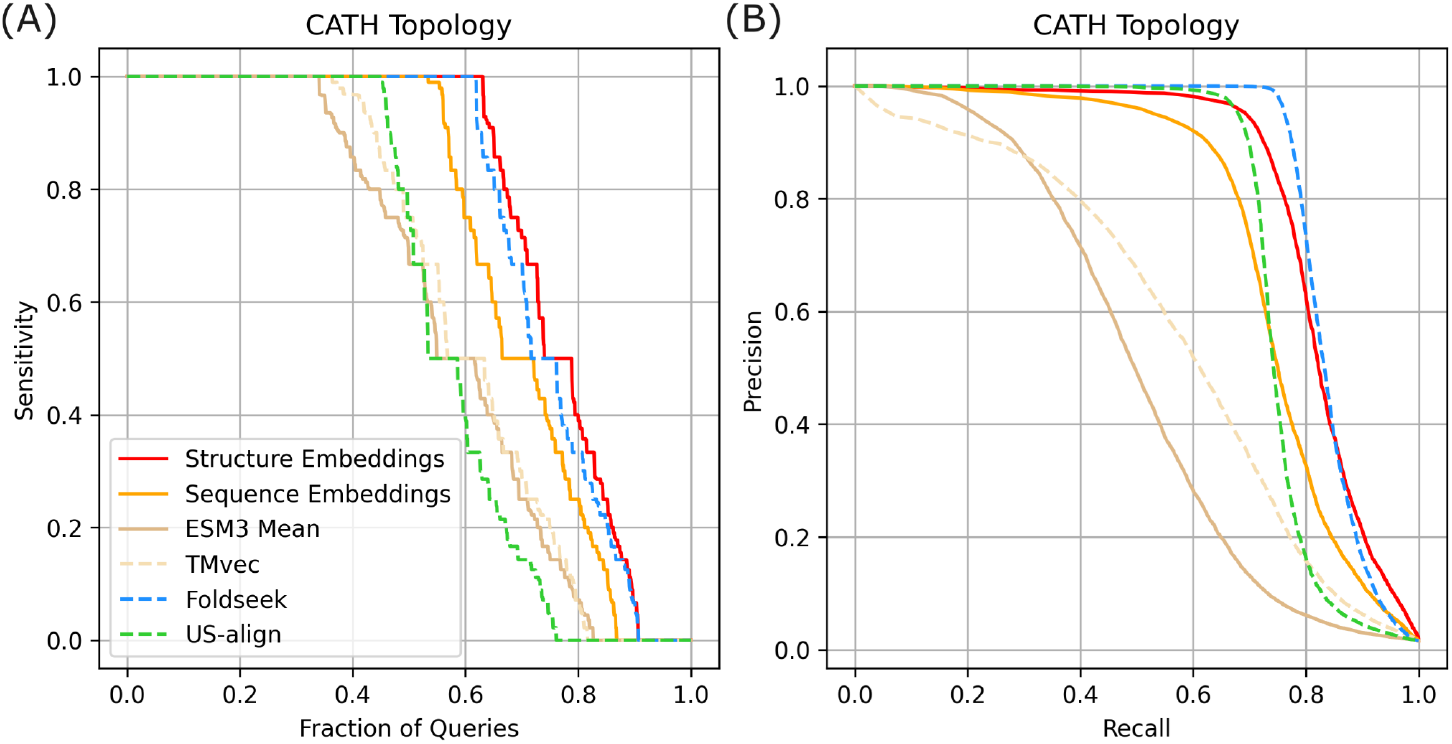
Sensitivity and precision-recall performance for a non-redundant set of AlphaFold DB protein structure models. This dataset consists of multidomain proteins defined by the CATH classification. Each protein model was compared against all others in the dataset, and results were ranked in descending order of predicted similarity score. True positives (TPs) were defined as proteins with the same combination of CATH domain topologies, and false positives (FPs) as proteins in which all domains belonged to different CATH architectures. (A) Sensitivity curves across methods. Sensitivity for each structure model was calculated as the number of TPs retrieved before the first FP; values are shown sorted from best to worst as a function of the fraction of queries. (B) Precision–recall curves across methods.

Figure 4A shows sensitivity values as a function of query fraction. Our embedding model and Foldseek performed comparably, achieving perfect sensitivity for 64% and 63% of queries, respectively. Precision–recall analysis (Figure 4B) revealed similar curves, with AUC values of 0.82 for the embedding model and 0.84 for Foldseek. As in previous benchmarks, sequence-based embeddings underperformed relative to structure-based approaches (precision–recall AUC of 0.75 for sequence-based embeddings and 0.58 for TMvec). Averaging ESM3 residue embeddings again resulted in reduced performance compared to adding the aggregator module in the embedding generation (precision–recall AUC = 0.50).

By contrast, US-align did not reproduce the strong performance observed on the domain benchmark. In this benchmark, it achieved a precision–recall AUC of 0.75, and its sensitivity was comparable to that of sequence-based embedding methods (Figure 4). This behavior is expected given the rigid superposition and global similarity that US-align is based on, which limit its ability to detect local domain-level similarities. Overall, these results confirm that our embedding model generalizes effectively to CSMs, providing robust performance in scenarios where global superposition methods may struggle.

### 3.4 Assembly Benchmark

Structural similarity search at the assembly level is a valuable capability that only a few open-access resources currently provide (Kim *et al*., 2025). The primary goal of this benchmark was to evaluate the performance of our embedding model in identifying multimeric assemblies with similar quaternary structures. In addition, this analysis demonstrates that the latent features learned by our embedding model extend beyond single protein chains or domains to capture assembly-level organization.

For this benchmark, we used assembly structures from the AS50M dataset and evaluated sensitivity to the first FP and precision–recall performance using the same methodology described in the full-chain benchmark (Section 3.2). TPs were defined using TM-score thresholds of 0.6, 0.7, and 0.8, while FPs were pairs with TM-scores below 0.5. Figure 5 presents the results for the strictest threshold (TM-score ≥ 0.8), with the additional results available in Supplementary Material Figure S7 and Table S5. As in the full-chain benchmark, US-align served as the ground truth and therefore achieved a perfect classification. Foldseek-MM achieved the best performance, with 84% of assemblies reaching perfect sensitivity (Figure 5A) and the highest precision–recall (AUC 0.97; Figure 5B). Our embedding model performed behind Foldseek-MM, attaining 100% of sensitivity for 77% of the queries and a precision–recall AUC of 0.92. As observed in the other benchmarks, sequence-based embeddings performed behind our structural-based model with perfect sensitivity for 68% of the assemblies and a precision–recall AUC of 0.89.

**Figure 5.**
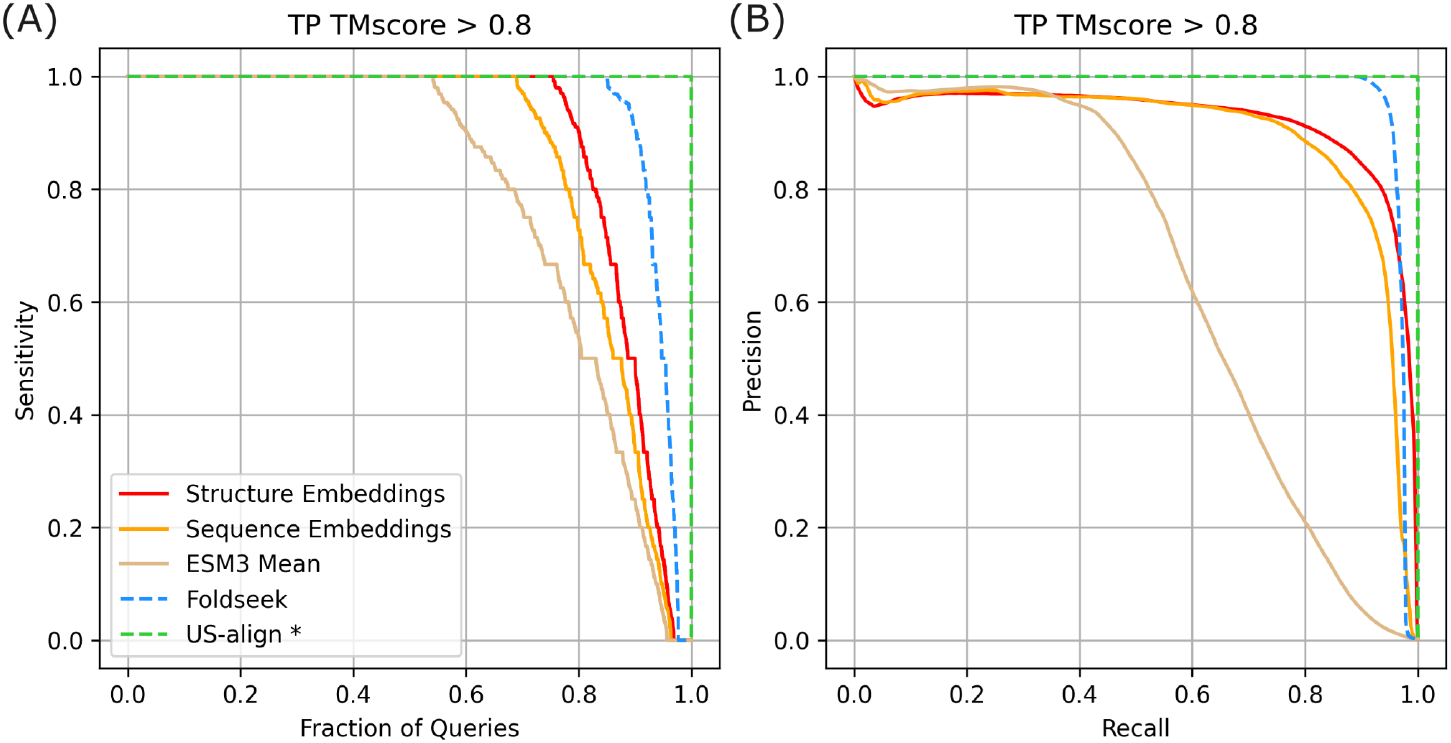
Sensitivity and precision-recall performance for a random subset of the 3D Complex DB assemblies. Each assembly was compared against all others in the dataset, and results were ranked in descending order of predicted similarity score. True positives (TPs) were defined as matches with TM-score > 0.8, and false positives (FPs) as matches with TM-score < 0.5. (A) Sensitivity curves across methods. Sensitivity for each assembly was calculated as the number of TPs retrieved before the first FP; values are shown sorted from best to worst as a function of the fraction of queries. (B) Precision–recall curves across methods. US-align* shows perfect performance because it was used to compute the TM-score ground truth.

To better understand the limitations of our approach in assembly-level searches, we examined high-scoring FPs produced by our embedding model. Many of these cases involved assembly pairs with highly similar monomeric subunits (high TM-scores between monomers) but distinct overall conformations (TM-score < 0.5 for the assemblies). Supplementary Material Figure S8 shows the TM-score distribution for the monomeric subunits of the assembly benchmark for the first 10,000 FP pairs, where more than 80% of the cases have a TM-score > 0.8.

This outcome reflects a known limitation of our methodology: residue embeddings are computed independently at the chain level and then concatenated before being processed by the aggregator module to generate the final assembly embedding. Consequently, if identical subunits (i.e., with the same sequence and 3D structure) assemble in different arrangements in different assemblies, the model computes the same embedding for all of them. Likewise, assemblies composed of similar monomers—though not strictly identical—but arranged in distinct quaternary structures may yield closely related embeddings.

Despite this limitation, our embedding approach achieved competitive results relative to Foldseek-MM. Furthermore, explicitly recognizing these constraints helps users interpret potential false positives and provides a basis for improving future versions of the method.

### 3.5 Scaling and Runtime

To assess the scalability and efficiency of our structural similarity search based on embeddings and vector databases, we evaluated its runtime performance across two datasets that represent extensive catalogs of protein structures: (1) all macromolecular structures from RCSB.org, which includes approximately 2 million chains from experimentally determined structures (from PDB) and CSMs (from AlphaFold DB and the ModelArchive), and (2) the 214 million predicted structures available from AlphaFold DB.

Both datasets were stored in independent Milvus database instances optimized for large-scale and fast vector retrieval. Chains from RCSB.org were indexed using the HNSW algorithm, and the database engine runs on a 32GB RAM, 8-CPU instance. The much larger AlphaFold DB dataset was indexed using DiskANN, a disk-based ANN method designed for large-scale retrieval with minimal memory overhead running on a 128GB RAM, 8-CPU instance. Search performance was evaluated by measuring query retrieval times as a function of the number of retrieved results. These results do not include the computational cost associated with generating the query embedding, which represents a fixed one-off computational cost. Due to the nature of ANN algorithms, retrieving a larger number of neighbors (results) requires additional graph index exploration, leading to increased query times. Table 1 presents the average retrieval times for multiple searches, considering varying numbers of results returned per query and dataset (RCSB.org chains and AlphaFold DB).

**Table 1.** Query Performance Benchmark for Structural Similarity Search. Average retrieval time for 100 queries. Times are given in seconds. Retrieval time variances for RCSB.org chains were negligible.

| Number of Results | RCSB.org chains (2M) | AlphaFold DB (214M) |
| --- | --- | --- |
| 10 | 0.003 | 0.5 ± 0.1 |
| 100 | 0.004 | 1.55 ± 0.15 |
| 1000 | 0.02 | 10.6 ± 0.2 |

As shown in Table 1, searches across all RCSB.org chains are essentially instantaneous, with average retrieval times of 0.02 seconds when returning 1000 results. Searches across AlphaFold DB exhibit longer runtimes, ranging from 0.5 seconds (10 results) to 10.6 seconds (1000 results). Despite this difference, the DiskANN-based approach remains highly efficient given the scale of the AlphaFold DB dataset, offering real-time structural retrieval capabilities across the equivalent of many thousand proteomes in their entirety.

Importantly, vector databases and ANN algorithms are an active field of research, so it is reasonable to expect more efficient systems to emerge (Aguerrebere *et al*., 2023) to further improve the scalability and enable applicability to even larger datasets. It is also noteworthy that our current implementation runs on a single instance, and available vector database engines (e.g., Milvus, Elasticsearch, MongoDB) offer horizontal scalability capabilities, thus already allowing scale-ups to whole computer clusters.

## 4 Conclusion

In this work, we introduced a new structural similarity search strategy based on 3D structure embeddings and vector databases, capable of scaling to vast repositories of macromolecular structures (including both experimentally determined models and CSMs). Trained to predict TM-scores between single-domain structures, our approach demonstrated strong generalization beyond the domain level to full-length protein chains and even multi-protein assemblies. The embedding model achieved sensitivity levels comparable to those of state-of-the-art methods, while providing a highly scalable and computationally efficient alternative to traditional structural alignment techniques. Extensive benchmarking confirmed that our embedding model effectively captures structural similarities at multiple levels of granularity, aligning well with established protein domain classification databases. Moreover, its ability to retrieve structurally similar proteins and assemblies with high sensitivity highlights its potential for functional annotation, remote homology detection, and large-scale structural analysis. By integrating our method with a vector database infrastructure optimized for real-time retrieval, we enable efficient and scalable structural searches across expanding structural datasets, including AlphaFold DB with >214 million predicted structures, representing a significant step towards real-time, large-scale structural similarity searches for computational structural biology.

## Acknowledgments

RCSB PDB Core Operations are funded by the US National Science Foundation (DBI-2321666, P.I.: S.K. Burley), the US Department of Energy (DE-SC0019749, P.I.: S.K. Burley), and the National Cancer Institute, National Institute of Allergy and Infectious Diseases, and National Institute of General Medical Sciences of the National Institutes of Health under grant R01GM157729 (P.I.: S.K. Burley). We would also like to acknowledge the NSF ACCESS program that provided credits for access to the Expanse supercomputer at the San Diego Supercomputer Center. Additionally, access to the DOE NERSC supercomputer was granted as part of DE-SC0019749.

