## Supplementary Material for "Multi-scale structural similarity embedding search across entire proteomes"



Code availability: https://github.com/bioinsilico/rcsb-embedding-search

Code DOI: https://doi.org/10.6084/m9.figshare.30006019.v1

Data availability: https://doi.org/10.6084/m9.figshare.30005845.v1

Web server: http://embedding-search.rcsb.org/

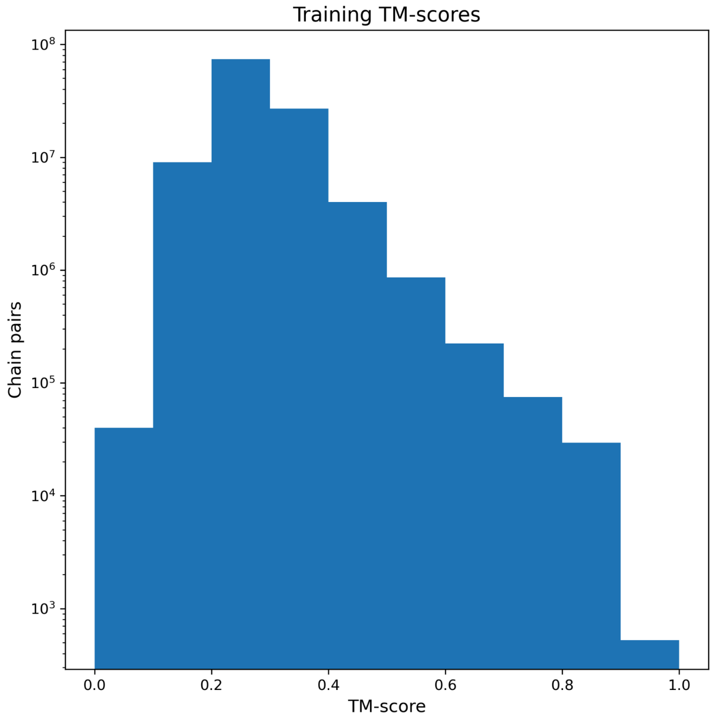


**Figure S1**. **TM-score distribution for the SCOPe40 v2.08 training pairs**. Histogram of TM-scores for domain pairs (>115M) used in training, illustrating an imbalanced distribution. The vast majority (>95%) of TM-scores are below 0.4, while fewer than 0.05% exceed 0.8.

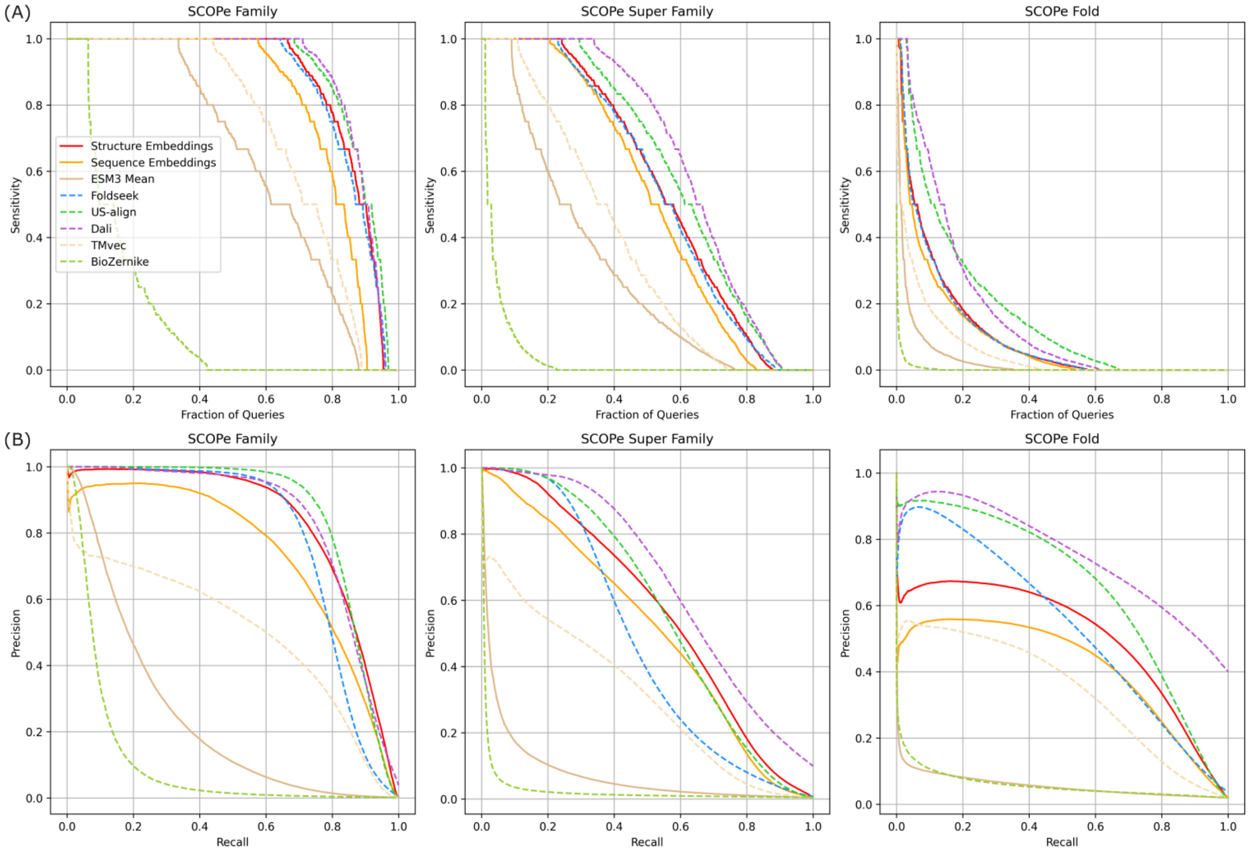


**Figure S2**. **Sensitivity and precision-recall performance on SCOPe40 v2.01 domain structures.** Each domain was compared against all others in the dataset, and results were ranked in descending order of predicted similarity score. True positives (TPs) were defined as matches within the same family (left), within the same superfamily but different families (middle), and within the same fold but different superfamilies (right). False positives (FPs) were defined as matches between different folds. (A) Sensitivity curves across methods. Sensitivity for each domain was calculated as the number of TPs retrieved before the first FP; values are shown sorted from best to worst as a function of the fraction of queries. (B) Precision–recall curves across methods.

| | Family | Superfamily | Fold |
|---|---|---|---|
| Structure-embedding | 0.83 | 0.57 | 0.51 |
| Sequence-embedding | 0.73 | 0.51 | 0.41 |
| ESM3 mean | 0.24 | 0.07 | 0.06 |
| Foldseek | 0.78 | 0.48 | 0.53 |
| US-align | 0.86 | 0.58 | 0.65 |
| Dali | 0.84 | 0.66 | 0.75 |
| TMvec | 0.50 | 0.31 | 0.34 |
| BioZernike | 0.10 | 0.03 | 0.06 |

**Table S1**. **Precision-recall AUC performance on SCOPe40 v2.01 domain structures**. Each domain was compared against all others in the dataset, and results were ranked in descending order of predicted similarity score. True positives were defined as matches within the same family (Family column), within the same superfamily but different families (Superfamily column), and within the same fold but different superfamilies (Fold column). False positives were defined as matches between different folds.

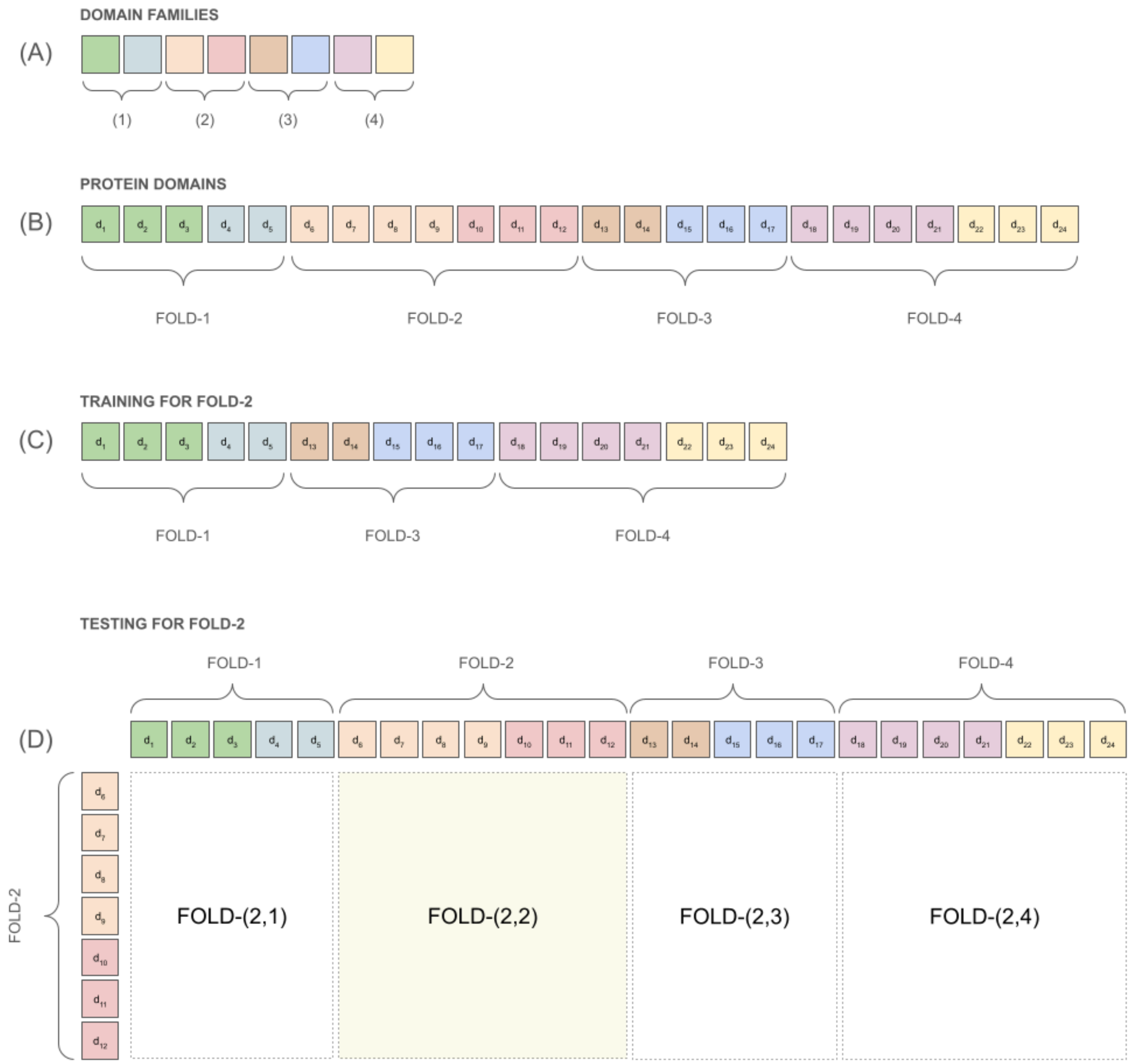


**Figure S3**. **SCOPe40 v2.01 fold-cross schema representation used for training and testing**. (A) Domain dataset containing eight families, grouped into four sets of two families each. (B) The dataset is divided into four folds, each fold including only domains from one of the family sets. (C-D) Example of cross-validation: domains in Fold-2 are held out for testing (D), while domains in Folds 1, 3, and 4 are used for training (C). In (D), the domain families assigned to Fold-2 are unseen during training. However, Fold-2 domains are compared against the entire dataset, so only domain pairs within Fold-(2,2) have both members excluded from training. In contrast, for Fold-(2,{1,3,4}), one domain of the pair was included in training. This cross-validation scheme ensures that at least one member of each test pair was not part of the training set.

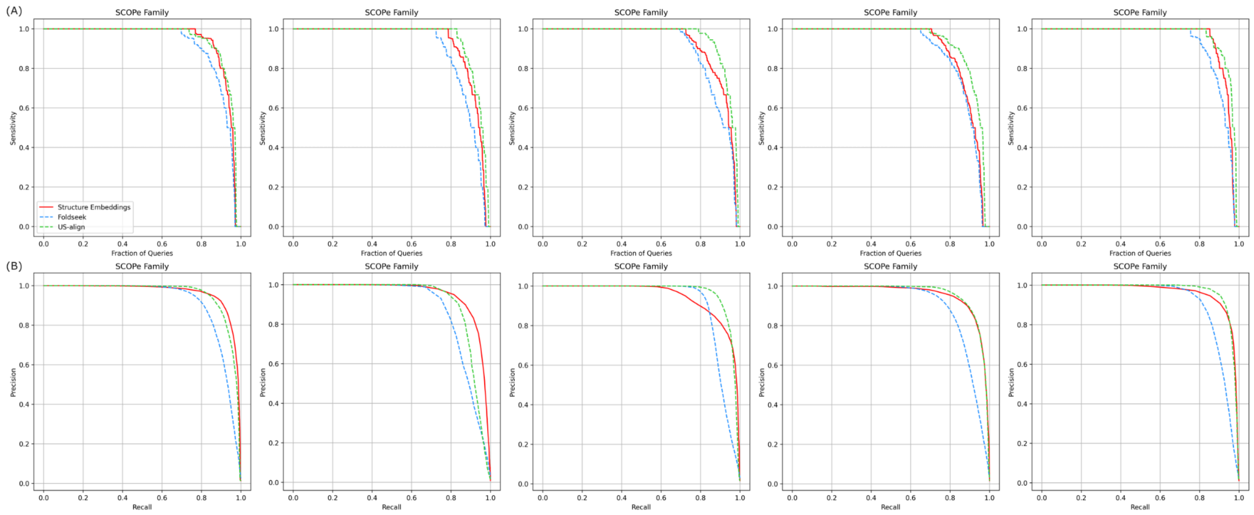


**Figure S4**. **Sensitivity and precision-recall results on SCOPe40 v2.01 held-out protein domains.** Evaluation of sensitivity and precision-recall curves on 5 random samples (different columns) of the SCOPe40 v2.01 domains. 5% of the different domain superfamilies were randomly selected, and all the domains belonging to these superfamilies were held out for testing. The remaining set of domains was used for training. The remaining domains were used for training. During testing, each held-out domain was compared against all others in the testing set. True positives (TPs) were defined as matches within the same family, and false positives (FPs) were defined as matches between different folds. (A) Sensitivity curves across methods. Sensitivity for each domain was calculated as the number of TPs retrieved before the first FP; values are shown sorted from best to worst as a function of the fraction of queries. (B) Precision–recall curves across methods.

| | Sample 1 | Sample 2 | Sample 3 | Sample 4 | Sample 5 |
|---|---|---|---|---|---|
| Structure-embedding | 0.96 | 0.94 | 0.93 | 0.95 | 0.95 |
| Foldseek | 0.91 | 0.89 | 0.91 | 0.90 | 0.91 |
| US-align | 0.95 | 0.91 | 0.96 | 0.96 | 0.97 |

**Table S2**. **Precision-recall AUC performance on SCOPe40 v2.01 held-out protein domains**. Evaluation was performed on six random samples of SCOPe40 v2.01 domains. For each sample, 5% of domain superfamilies were randomly selected, and all domains belonging to these superfamilies were withheld for testing. The remaining domains were used for training. The remaining domains were used for training. During testing, each held-out domain was compared against all others in the testing set. True positives were defined as matches within the same family, and false positives as matches between different folds.

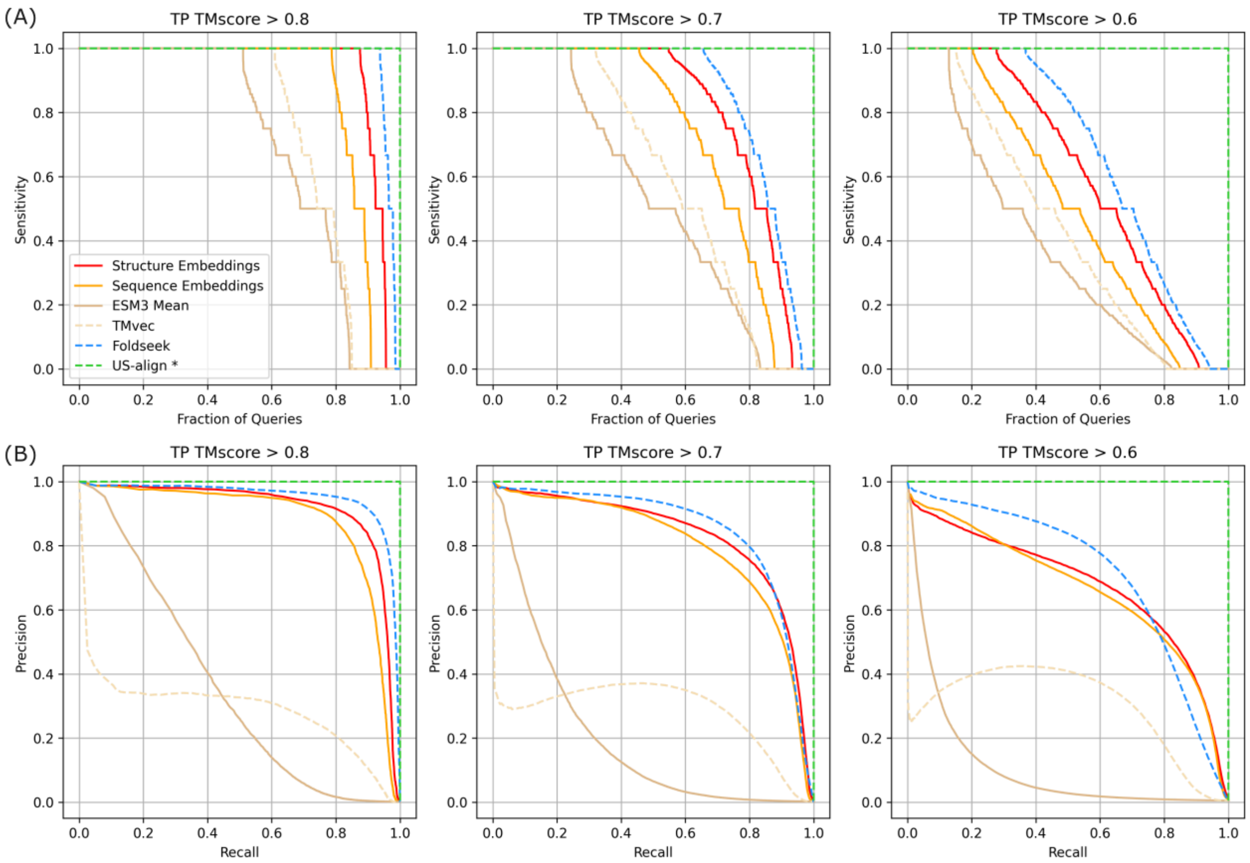


**Figure S5**. **Sensitivity and precision-recall performance for a non-redundant set of PDB protein chain structures**. Each protein chain was compared against all others in the dataset, and results were ranked in descending order of predicted similarity score. True positives (TPs) were defined as matches with TM-score > 0.8 (left), TM-score > 0.7 (middle), and TM-score > 0.6 (right), and false positives (FPs) as matches with TM-score < 0.5. (A) Sensitivity curves across methods. Sensitivity for each chain was calculated as the number of TPs retrieved before the first FP; values are shown sorted from best to worst as a function of the fraction of queries. (B) Precision–recall curves across methods. *US-align* shows perfect performance because it was used to compute the TM-score ground truth.

| | TM-score > 0.8 | TM-score > 0.7 | TM-score > 0.6 |
|---|---|---|---|
| Structure-embedding | 0.91 | 0.83 | 0.68 |
| Sequence-embedding | 0.88 | 0.80 | 0.67 |
| ESM3 Mean | 0.35 | 0.20 | 0.11 |
| TMvec | 0.29 | 0.28 | 0.30 |
| Foldseek | 0.94 | 0.85 | 0.71 |
| US-align | 1.0* | 1.0* | 1.0* |

**Table S3**. **Precision-recall AUC performance for a non-redundant set of PDB proteins**. Each protein chain was compared against all others in the dataset, and results were ranked in descending order of predicted similarity score. True positives were defined as matches with TM-score > 0.8, TM-score > 0.7, or TM-score > 0.6, and false positives as matches with TM-score < 0.5. US-align shows perfect AUC values because it was used to compute the TM-score ground truth.

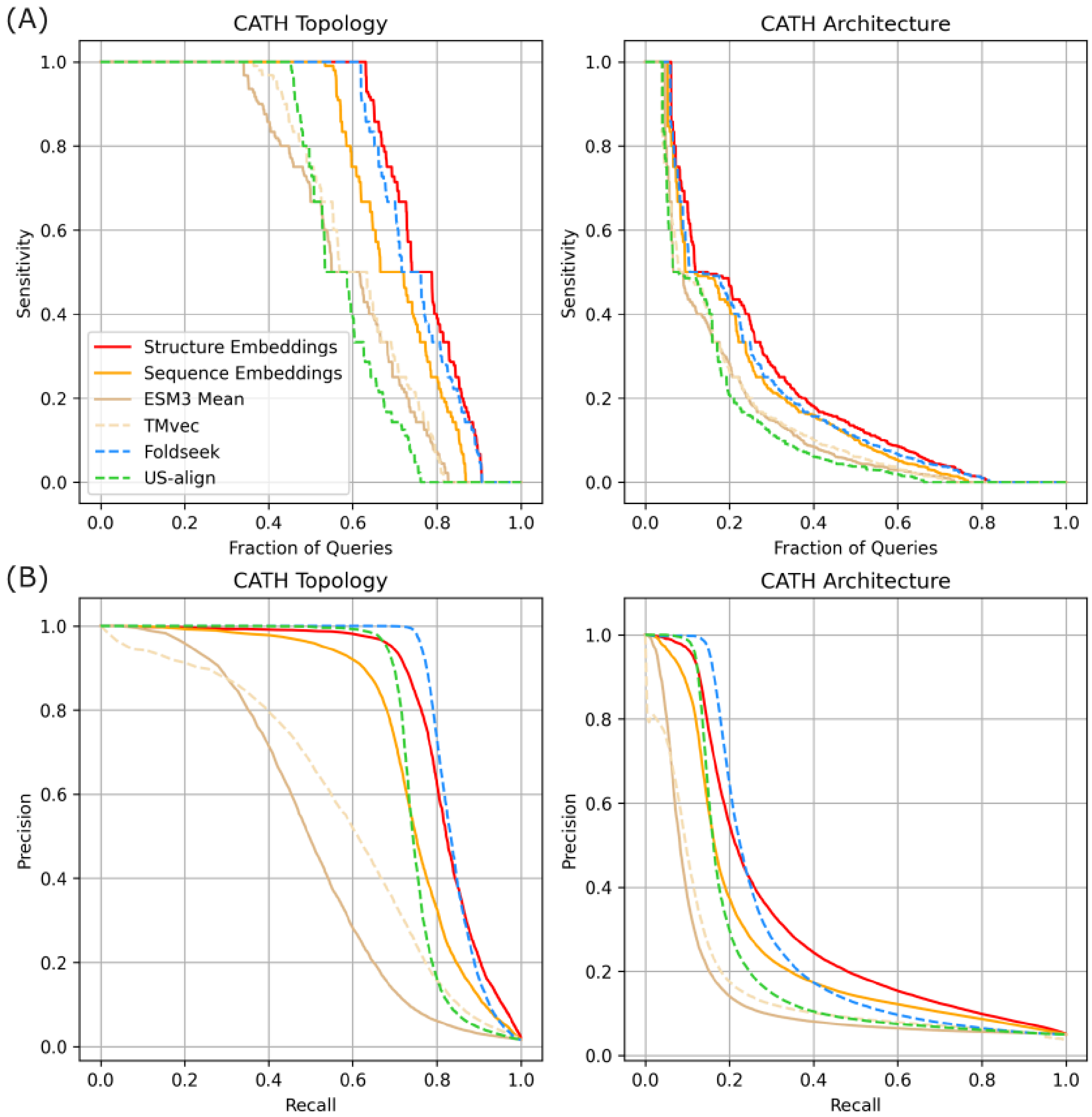


**Figure S6**. **Sensitivity and precision-recall performance for a non-redundant set of AlphaFold DB protein structure models**. The dataset consists of multidomain proteins defined by the CATH classification. Each protein model was compared against all others in the dataset, and results were ranked in descending order of predicted similarity score. True positives (TPs) were defined as proteins with the same combination of CATH domain topologies (left) or the same combination of CATH architectures (right), and false positives (FPs) as proteins in which all domains belonged to different CATH architectures. (A) Sensitivity curves across methods. Sensitivity for each structure model was calculated as the number of TPs retrieved before the first FP; values are shown sorted from best to worst as a function of the fraction of queries. (B) Precision–recall curves across methods.

| | Topology | Architecture |
|---|---|---|
| Structure-embedding | 0.82 | 0.33 |
| Sequence-embedding | 0.75 | 0.27 |
| ESM3 Mean | 0.50 | 0.15 |
| TMvec | 0.58 | 0.17 |
| Foldseek | 0.84 | 0.31 |
| US-align | 0.75 | 0.24 |

**Table S4**. **Precision-recall AUC performance for a non-redundant set of AlphaFold DB models**. The dataset consists of multidomain proteins defined by the CATH classification. Each protein model was compared against all others in the dataset, and results were ranked in descending order of predicted similarity score. True positives were defined as proteins with the same combination of CATH domain topologies (Topology column) or the same combination of CATH architectures (Architecture column), and false positives as proteins in which all domains belonged to different CATH architectures.

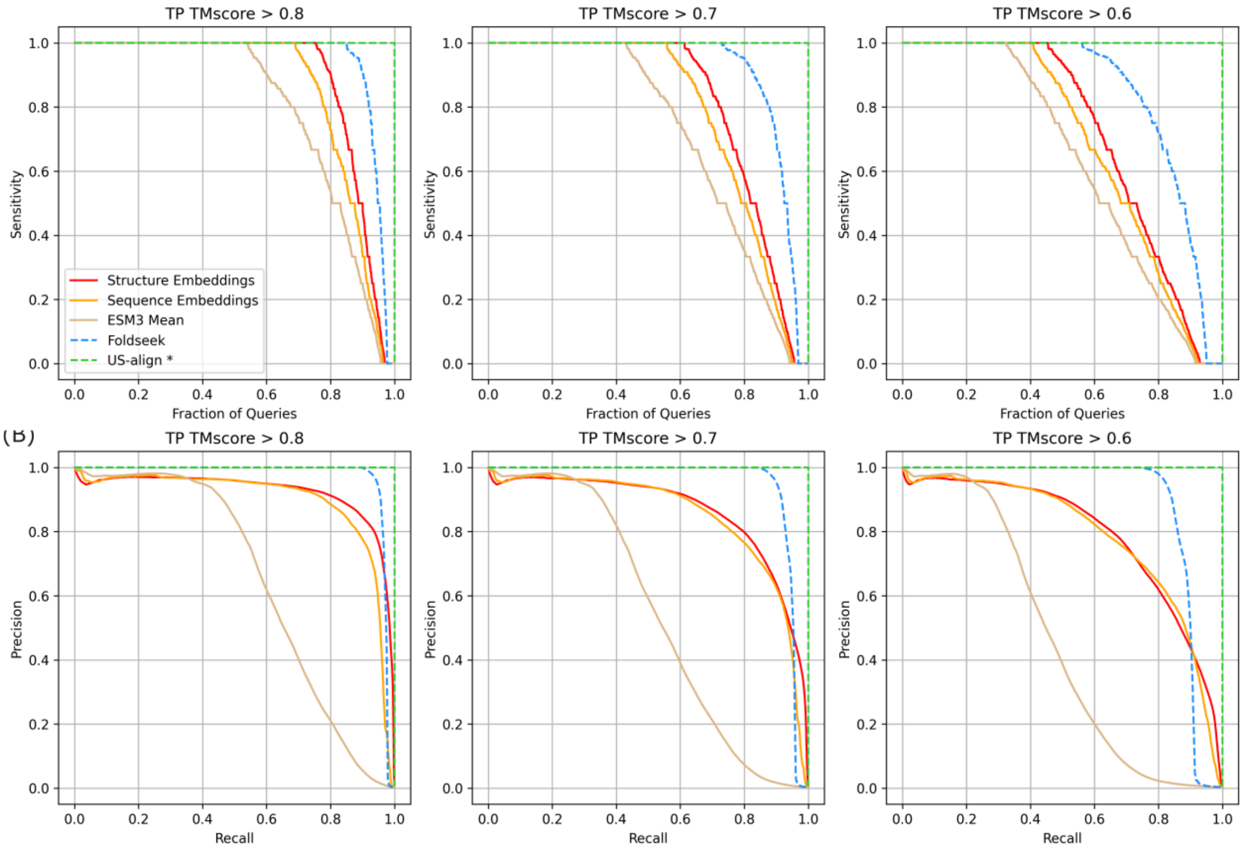


**Figure S7**. **Sensitivity and precision-recall results for a random subset of the 3D Complex DB assemblies**. Each assembly was compared against all others in the dataset, and results were ranked in descending order of predicted similarity score. True positives (TPs) were defined as matches with TM-score > 0.8 (left), TM-score > 0.7 (middle), and TM-score > 0.6 (right), and false positives (FPs) as matches with TM-score < 0.5. (A) Sensitivity curves across methods. Sensitivity for each assembly was calculated as the number of TPs retrieved before the first FP; values are shown sorted from best to worst as a function of the fraction of queries. (B) Precision–recall curves across methods. *US-align* shows perfect performance because it was used to compute the TM-score ground truth.

| | TM-score > 0.8 | TM-score > 0.7 | TM-score > 0.6 |
|---|---|---|---|
| Structure-embedding | 0.92 | 0.86 | 0.79 |
| Sequnce-embedding | 0.89 | 0.84 | 0.79 |
| ESM3 Mean | 0.65 | 0.55 | 0.46 |
| Foldseek | 0.97 | 0.94 | 0.88 |
| US-align | 1.0* | 1.0* | 1.0* |

**Table S5**. **Precision-recall AUC results for a non-redundant set of assemblies**. Each assembly was compared against all others in the dataset, and results were ranked in descending order of predicted similarity score. True positives were defined as matches with TM-score > 0.8, TM-score > 0.7, or TM-score > 0.6, and false positives as matches with TM-score < 0.5.

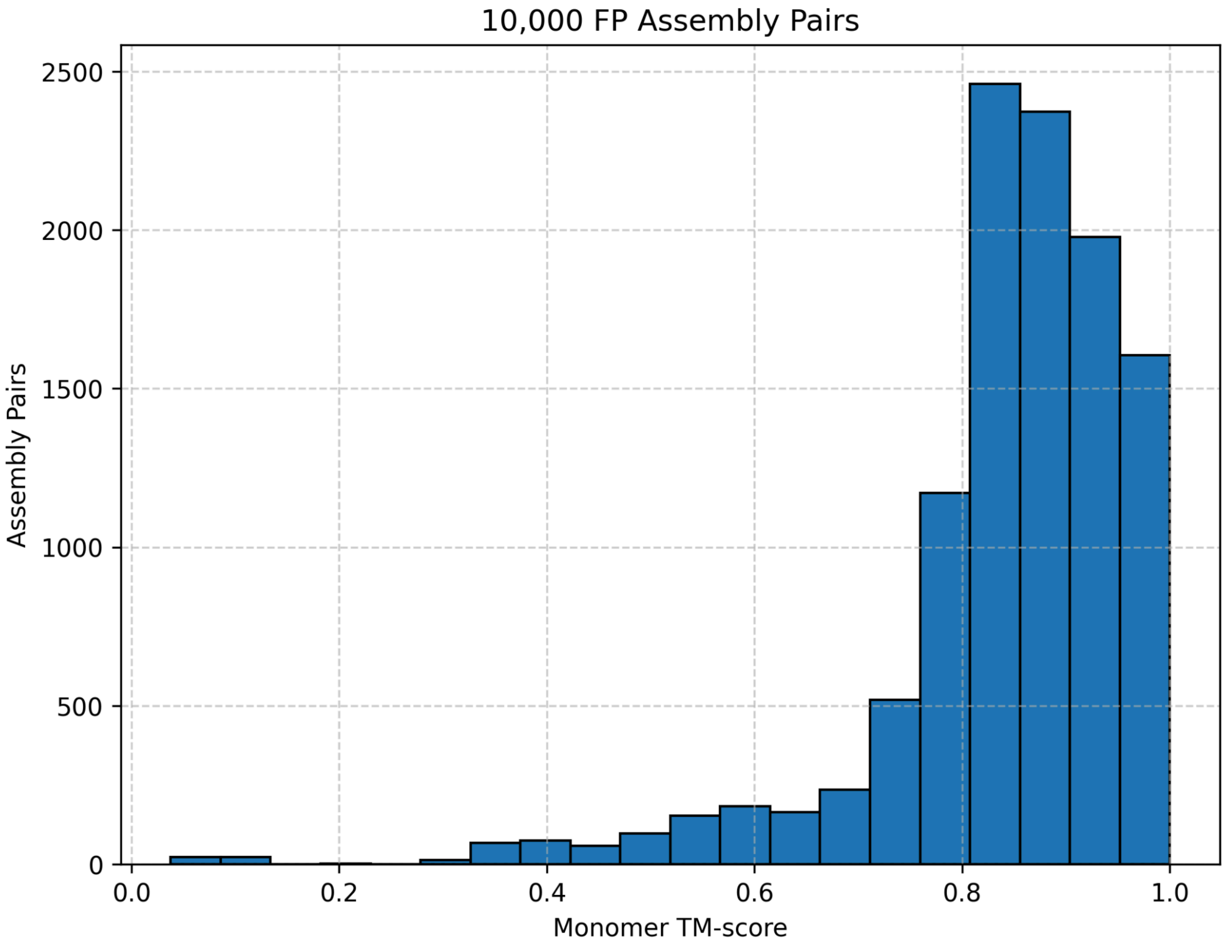


**Figure S8**. **FP assembly pairs TM-score distribution**. Distribution of TM-scores between the monomeric subunits of the 10,000 highest-scoring FP pairs in the homomeric assembly benchmark. More than 80% of these pairs have a TM-score > 0.8.
